# Long-term effects of ocean acidification and its interaction with warming on calcifying organisms and their associated microbiome: bryozoans as emerging sentinels of global change?

**DOI:** 10.1101/2024.06.28.601120

**Authors:** Blanca Figuerola, Pol Capdevila, Marc Cerdà-Domènech, Joaquim Garrabou, Alice Mirasole, Pol Bassols, Javier del Campo, Núria Teixidó

## Abstract

Global ocean warming and acidification are two of the major threats to many marine calcifying habitat-forming species, potentially affecting entire ecosystems. Consequently, the need for a better understanding and predicting the response of marine calcifiers has never been more pressing. Paradoxically, the individual and combined long-term effects of these stressors on bryozoans have remained largely unexplored, despite their great abundance and diversity globally. Here, we first evaluate the changes in skeletal structure and mineralogy, and the associated microbiome composition on the populations of *Pentapora ottomuelleriana* (encrusting) and *Myriapora truncata* (erect) bryozoan species living inside and outside a volcanic CO_2_ vent in Ischia Island. We then examine the effects of a long-term exposure to elevated *p*CO_2_ and its combined effects of ocean warming on the proportion of cover of populations of the encrusting species through time after summer. Both bryozoan species show indicators of acclimatization by adjusting skeletal properties and having stable microbial communities under acidification conditions. However, we document novel patterns about microbiome shifts in response to future ocean acidification in bryozoans for the first time. Microbial genera known to have essential functions to the host such as biosynthesis of defense compounds or thermal protection were depleted at the acidified site, which suggest early warnings of potential deterioration of bryozoan health under near future ocean conditions. The proportion of cover of the encrusting species also decreased from 2016 to 2020 in both studied sites, with faster declines at the acidified ones. Our model suggests that the increasing seawater temperature drove a decline in the bryozoan cover although the combined effects with acidification accelerated its mortality rates in the CO_2_ vent. More multidisciplinary research combining both environmental stressors on a wider range of calcifying species is needed to better understand the adaptive capacity of the holobiont to a changing environment.

**Graphical abstract:** 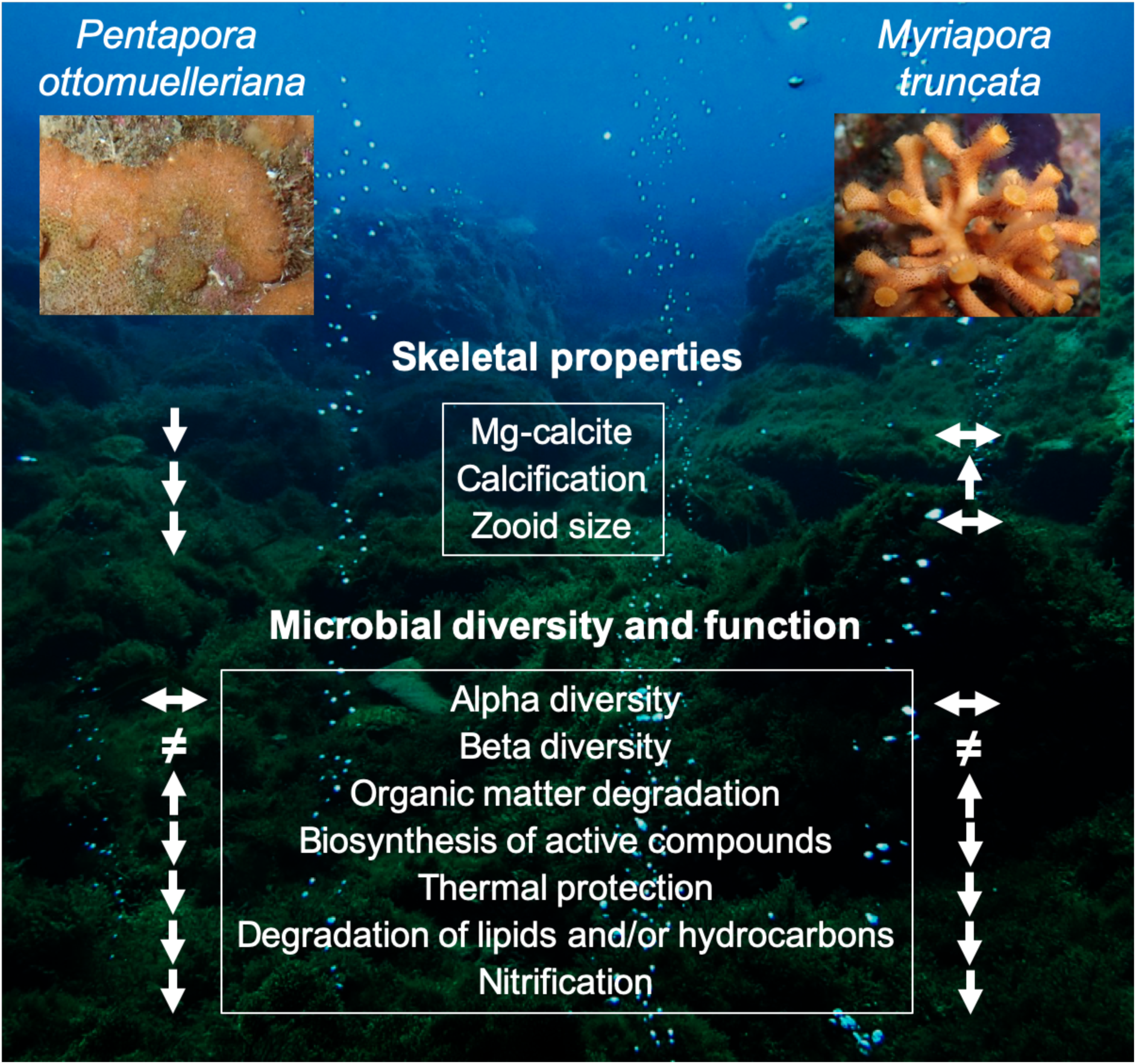

## 1. Introduction

Global environmental change brings multifaceted impacts to species worldwide (IPCC, 2022; Sage, 2020). In the marine realm, two of the major threats are ocean warming and ocean acidification. The combination of these two facets of global environmental change, in particular, threatens marine calcifiers, which build carbonate structures and often are habitat-forming species, such as corals (Carbonne et al., 2021; Cornwall et al., 2021). Some calcifying species may be impacted with increased skeletons/shells fragility and/or extra energetic cost for maintaining their carbonate structures. These effects are particularly strong in those species made from more soluble mineral phases such as aragonite or high Mg-calcite (Fabry, 2008; Figuerola et al., 2021). Therefore, even in the most optimistic scenarios, temperatures and acidification are going to increase and may compromise many calcifying species at multiple levels, resulting in shifts in marine biodiversity and functioning (Orr et al., 2005; Teixidó et al., 2018), with negative effects for other species and ecosystem services (*e.g.*, pharmacological products). A better understanding and predicting the response of marine calcifiers is thus more urgent than ever.

Assessing the responses of marine calcifiers to the multifaceted threats of climate change is challenging, as co-occurring environmental drivers can cause complex responses. Specifically, both drivers (seawater temperature and pH) can have synergistic or antagonistic effects – the cumulative effects of both drivers are larger than or smaller than the sum of their individual effects, respectively –, or even no effects on calcification (Cross et al., 2019; Prada et al., 2017). Phenotypic plasticity (*e.g.*, physiological, developmental, mineralogical and structural morphological adjustments) will thus determine the ability of calcifiers to cope with these rapid environmental changes. For instance, growing evidence reveals that a range of calcifying species can acclimatize to rapid environmental changes and potentially lead to local adaptation (*e.g.*, calcifying species living in coastal zones with temporal fluctuations in temperature and/or pH) (Gaitán-Espitia et al., 2017; Li et al., 2018; Pespeni et al., 2013). However, it remains unknown if these species will be able to adapt and survive at the rate and magnitude of environmental change (Fox et al., 2019).

To properly understand how marine calcifiers will respond to near-future ocean conditions, it is also crucial to study their associated microbiome (assemblage of bacteria, archaea, microbial eukaryotes, and viruses). Microbes play an important role in the health and survival of the host (*e.g.*, recycling of nutrients and production of defensive compounds). Therefore, shifts in the microbiome composition are often considered detrimental to the hosts. For instance, changes in the microbiome can affect the health of marine calcifiers or even compromise their ability to adjust to the new environmental conditions (Leray et al., 2021; McDevitt-Irwin et al., 2017). However, microbiome can have the capacity to acclimatize faster than their hosts to changing conditions and may thus facilitate adaptive responses for their hosts (Torda et al., 2017). The few studies examining the interactive effects of increasing temperature and ocean acidification on their microbiome have already shown contrasting responses in different calcifying species (Webster et al., 2016), highlighting our limited understanding of the sensitivity of associated microbiome to near future climate scenarios.

Most current studies have been focused on short-term responses of calcifiers (*e.g.*, changes in calcification rates) to one stressor. These studies are mostly done through laboratory studies, while the combined and long-term effects of warming and acidification, through field experiments of natural populations, are scarce and taxonomically biased (Figuerola et al., 2021; Kroeker et al., 2013). However, long-term field experiments, combining these two major stressors and using a wider range of phyla and taxa with different skeletal mineralogies and biological traits, are key for understanding the adaptive ability of the holobiont (individual host and its microbiome) to near future ocean scenarios.

Bryozoans represent a good model species to study the complex impacts of global warming and acidification in marine calcifiers. This phylum of filter-feeding invertebrates are globally more speciose (c. 6000 extant species) (Bock and Gordon, 2013) and abundant (from poles to tropics and from the intertidal zone to the deep sea) (Figuerola et al., 2018, 2012; Rosso and Di Martino, 2016; Taylor and Tan, 2015) than other more well-studied calcifiers such as corals (Simpson and Jackson, 2022). Like corals, many bryozoan species are also habitat-forming organisms and important carbonate-producers that provide fundamental ecosystem services through protection from waves action (*e.g.*, Western Port bryozoan reefs), bioactive products, and tourism (Dutka et al., 2022; Figuerola and Avila, 2019; Wood et al., 2012). This phylum is thus well represented in the fossil record (Taylor, 2020) and represents a good model system for understanding ecological and evolutionary processes (Orr et al., 2022). In contrast to many invertebrates, bryozoans can secrete skeletons with either or both mineral forms (calcite and aragonite) of calcium carbonate and incorporate varying amounts of Mg in their calcite, which determine the degree of skeletal solubility and thus their potential vulnerability to ocean acidification (Smith, 2009). Their Mg content appears to be influenced by diverse environmental (*e.g.*, salinity, pH, temperature) and/or biological (*e.g.*, skeletal growth rate) factors on a local scale (Taylor, 2020). In contrast, at global scale, their skeletal Mg content is likely to be mainly influenced by seawater temperature (Figuerola et al., 2023). Therefore, they are suitable model holobionts to study the effects of global change. However, studies assessing short-term responses of bryozoan natural populations to elevated temperature and/or low pH stressors (Lombardi et al., 2011; Pagès-Escolà et al., 2018; Rodolfo-Metalpa et al., 2010) and the characterization of its associated microbiome (Boscaro et al., 2022; Bourne et al., 2013; Li et al., 2019; Patin et al., 2019) are limited and long-term responses to both stressors have not previously been assessed.

The Ischia Island (Naples, Italy) presents shallow underwater volcanic CO_2_ vents, which cause local acidification of seawater at pH levels predicted by 2100, being thus natural analogues to explore long-term responses of natural populations of marine calcifiers and their associated microbes to future ocean acidification conditions (Biagi et al., 2020; Carbonne et al., 2021; Teixidó et al., 2020). These CO_2_ vent systems are also particularly valuable for studying the combined effects of warming and acidification as they are located in the Mediterranean Sea, which is warming 20% faster than the global average (Lionello and Scarascia, 2018).

In this study, we selected two bryozoan calcified species with contrasting life-history traits and skeletal properties, *Pentapora ottomuelleriana* (encrusting) and *Myriapora truncata* (erect) that naturally co-occur in volcanic CO_2_ vents. We first evaluated the changes in skeletal structure, mineralogy and the associated microbiome composition on populations of both species living inside and outside a volcanic CO_2_ vent in Ischia Island. We then examine the effects of long-term exposure to elevated *p*CO_2_ and its combined effects of ocean warming on the proportion of cover of the encrusting species through time after summer. We expected that both bryozoan species living in the volcanic CO_2_ vent will show indicators of adaptation or acclimatization to elevate *p*CO_2_ but showing different compensatory mechanisms (*e.g.*, mineralogical adjustments) to counter the low pH due to their different life-history traits and skeletal mineralogy (Poluzzi and Sartori, 1975; Rodolfo-Metalpa et al., 2010; Taylor et al., 2009). In addition, we expected that both species have distinct and stable microbial communities at the low pH site, given the microbiome’s ability to quickly acclimatize to environmental changes (Torda et al., 2017). We also hypothesized that the encrusting species would display signs of mortality due to low pH and the exceptional thermal conditions experienced in this region during summers as already occurred in different calcifying species, including bryozoans (Garrabou et al., 2022).

## 2. Material and methods

### 2.1 Selected species and study area

The study was conducted at two sites at the Island of Ischia (Gulf of Naples, Italy), where populations of *Pentapora ottomuelleriana* and *Myriapora truncata* naturally inhabit (Fig. 1A and B). *P. ottomuelleriana* is an encrusting species that grows faster than *M. truncata*, found on rocky bottoms, on *Posidonia oceanica* rhizomes, and on leaves, mainly between 0–50 m depth. In contrast, *M. truncata* is an erect branching bryozoan with a widespread occurrence in the Mediterranean Sea, inhabiting hard rocky habitats between 1–130 m depth (Cocito and Ferdeghini, 1998; Gautier, 1962; Gristina and Balduzzi, 1999; Pisano and Boyer, 1985).

**Fig. 1.**
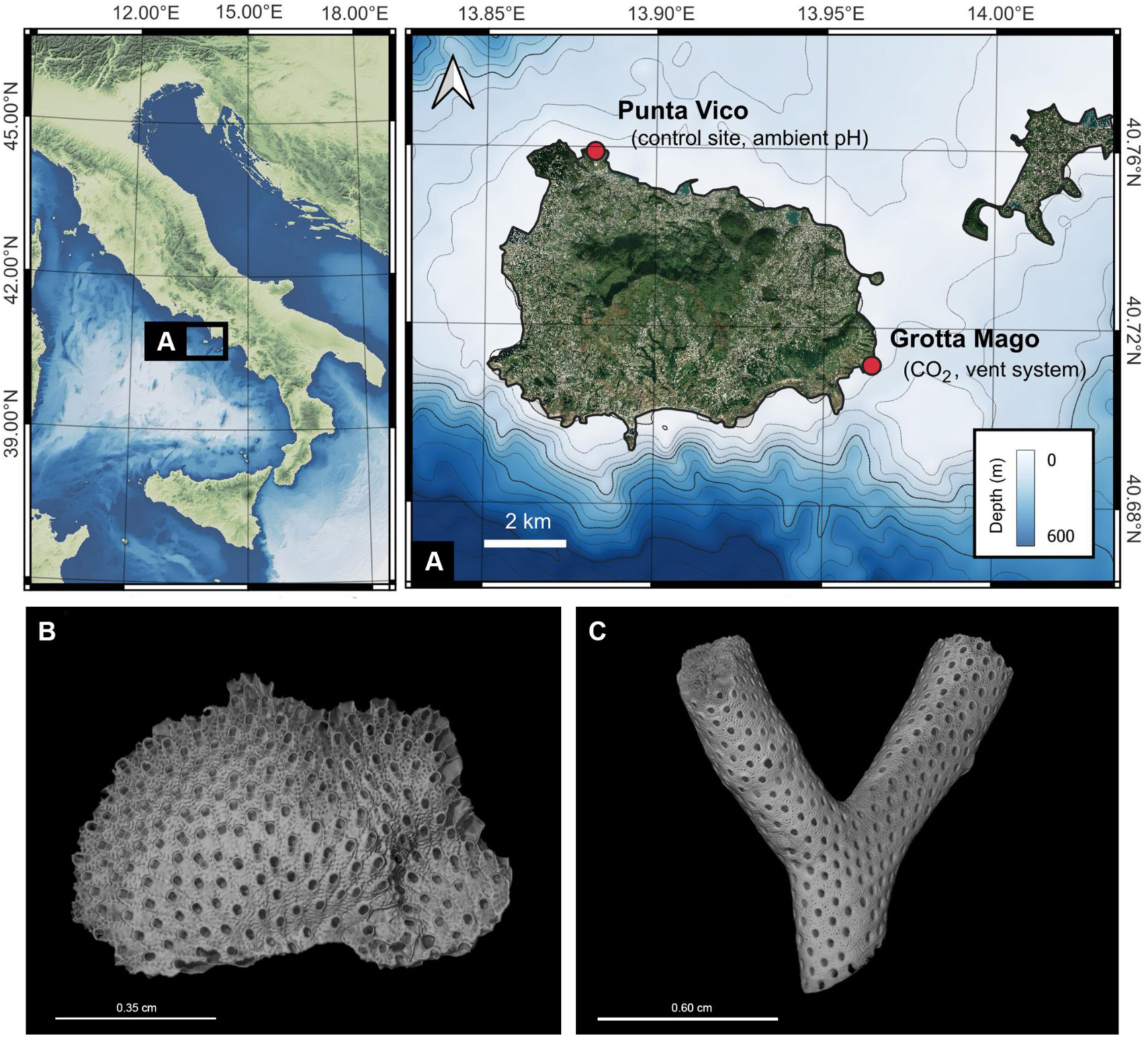
Study sites and target species. A. Map showing the location of the study sites along the coast of Ischia Island, Italy. B, C. *Pentapora ottomuelleriana* (left) and *Myriapora truncata* (right).

The study sites included an underwater volcanic CO_2_ vent and one reference site with ambient pH and no vent activity. The CO_2_ vent system is located at 5 m depth inside a semi-submerged cave of volcanic origin named Grotta del Mago (40°42′41.87″N, 13°57′51.06″E, hereafter CO_2_ vent). The cave has a main outer chamber (10 m wide x 30 m long), where the CO_2_ vent activity has increased over the last 50 years, with a more intense activity from 2014 (Cinelli et al., 1977; Teixidó et al., 2020). The reference site named Punta Vico is also a semi-submerged cave at the same depth which has a main chamber with the same dimensions (40°45′32.28″N,13°52′55.38″E, hereafter Ambient pH site). The present study was performed in the main chamber of both caves. The vent gas at the vent site was predominantly CO_2_, with undetectable levels of hydrogen sulfide (< 0.0002 %) (Teixidó et al. 2020). Water carbonate chemistry and *in situ* monitoring of seawater pH_T_ (pH on the total scale) at the CO_2_ vent site showed mean pH_T_ = 7.88 and 7.65 with 14% and 56% of the pH_T_ measurements were below 7.8 in June and September at 2 m depth, respectively. These values below 7.8 correspond to the projected average global sea surface pH value for the year 2100 with the high-CO_2_ emission scenario SSP5 RCP8.5 (Gattuso et al., 2015). Calcite (Ωc) and aragonite (Ωa) saturation states were 2.2 and 1.44 in June and 1.68 and 1.58 in September. The Ambient pH site showed pH_T_ = 8.05, Ωc = 5.06, and Ωa= 3.86 (for more information see Teixidó et al. 2020).

### 2.2 Temperature time series

We deployed temperature data loggers (HOBO TidbiT v2, Onset) in both sites over a 6-year period (2016-2021) at 2 m depth that registered temperature every hour, following ambient seasonal fluctuations (Fig. S1). Additionally, coastal seawater temperature data from 5 m depth in Ischia between 2012 to 2023 were obtained from the T-MEDNet temperature database (www.t-mednet.org). Daily averages were obtained from hourly records.

### 2.3 Skeletal mineralogy, geochemistry and structure

#### 2.3.1 Mineralogical analyses

Three bryozoan colonies per species and site (n = 12) were collected at 2 m depth in May 2021 and preserved in solutions of 95% ethanol. We performed mineralogical analyses cutting the growing edge (2 × 2 mm^2^) following previously described methodology (Figuerola et al. 2015). The excised pieces are estimated to have been formed within a period of ∼1–6 months prior to the date collection as the estimated vertical growth of *M. truncata* and *P. ottomuelleriana* is 0.41 cm yr^−1^ and 3.5 cm yr^−1^ (estimated from another species belonging to the same genus, *P. fascialis*), respectively (Cocito and Ferdeghini, 1998; Gristina and Balduzzi, 1999).

The X-ray powder diffraction (XRD) measurements were carried out with a Bruker D8-A25 diffractometer, equipped with a Cu Kα (λ = 1.5418 Å) radiation source generated at a voltage of 30 kV and a current of 40 mA and a LynxEye position sensitive detector (PSD) at the Geosciencies Barcelona (GEO3BCN-CSIC). The scans were acquired between 25.40° and 32.70° in 2θ at 0.015° steps and time per step of 384s. Phase identification was performed with the DIFFRAC.EVA software together with the Powder Diffraction File PDF-2 and the Crystallography Open Database (COD). The mol% MgCO_3_ in calcite was calculated by measuring the position of the d104 peak, assuming a linear interpolation between CaCO_3_ and MgCO_3_ and recalibrated for the specific machine used.

The skeletons were categorized as intermediate-Mg calcite (IMC; 4–8 mol% MgCO_3_), and high-Mg calcite (HMC; >8 mol% molCO_3_) (Rucker and Carver 1969). The differences in the Mg content between sites were analyzed using a t-test.

#### 2.3.2 X-ray computed tomography

Three bryozoan colonies per species and site (n = 12) were collected at 2 m depth in May 2021, rinsed with distilled water and then air dried. The growing edges were scanned using micro X-ray computed tomography (Micro-CT) at the Service of Non-Destructive Techniques of the Museo Nacional de Ciencias Naturales (MNCN-CSIC). All scans were executed using an XTH 160 Nikon Metrology, with a molybdenum target. During the analysis, the X-ray source was operated at a voltage of 160 kV with current ranging from 70-85 μA. A total of 3015 projections were taken at an exposure of 354 msec. The resulting dataset had a voxel size of 11 μm^3^. The CT Pro 3D software was used for the reconstruction of the raw projection (Bartscher et al., 2016; Townsend et al., 2018).

The image analysis was achieved using Avizo® software of the CORELAB Laboratory from Universitat de Barcelona. Post-processing images workflow was developed to perform a morphometric analysis of samples (*i.e.*, carbonate structure thickness and zooid volumes, length and surface) (SI Material and Methods; Videos S1 and S2).

The thickness analysis results of the skeletons, and a thickness map image, which offered a visual depiction of the spatial distribution and variations in thickness across the sample, were presented in millimeters. For *P. ottomuelleriana*, the thickness analysis was conducted on a total section of the colony. In contrast, for *M. truncata*, the analysis was focused on the distal part of the branches to avoid potential biases.

### 2.4 Microbiome

#### 2.4.1 DNA extraction

Four bryozoan colonies per species and site (n = 16) were collected at 2 m depth in May 2021, placed in plastic bags filled with seawater from the collection point and transported in ice to the laboratory. Samples were then rinsed in milliQ water, preserved in 75% ethanol and frozen at –20°C until DNA extraction. A total of four pieces (∼ 1 mm^2^) were cut from the growing edge of each colony (n = 32) and transferred into a 1.5-mL Eppendorf tube. DNA was extracted from each sample using the DNeasy Blood & Tissue Kit (Qiagen, Hilden, Germany). All samples were stored at −20°C until further processing.

#### 2.4.2 DNA extraction from seawater samples

At each site, 1L of seawater was collected, transported in ice to the laboratory. Seawater samples were then filtered using 0.2-μm pore-size sterile membrane filters. Membrane filters were placed in plastic bags and stored at −20°C until DNA extraction. Microbiome DNA was then extracted using a Powersoil DNA extraction kit (Qiagen, Hilden, Germany). All samples were stored at −20°C until further processing.

#### 2.4.3 16S rRNA gene PCR amplification and sequencing

DNA concentrations from samples were measured with a Qubit fluorometer. Concentrations that were higher than 1ng/μ were sent to the Integrated Microbiome Resource facility at the Centre for Comparative Genomics and Evolutionary Bioinformatics at Dalhousie University, Canada for 16S rRNA gene amplification and sequencing. The V4 region of the prokaryotic 16S rRNA gene was PCR-amplified using the newly modified 515F/806RB primers with Illumina overhang adapter sequences. Sequencing was performed on Illuminia MiSeq platform.

#### 2.4.4 Bioinformatics and statistics

Raw sequencing data has been deposited on NCBI (BioProject: PRJNA1047010). Primers and adaptors used for sequencing were removed from paired-end reads using Cutadapt (Martin, 2011). Reads were then processed in R using the DADA2 package (Callahan et al., 2016). First, forward reads were truncated at 250 bp and reverse reads at 200 bp according to their quality profiles and filtered using the parameters maxN = 0 and maxEE = c(2,2). Paired-end reads were then merged and chimaeras removed and reads grouped into amplicon sequence variants (ASVs). Taxonomy was assigned to ASVs using the SILVA database as reference (Quast et al., 2013). The resulting ASV table and taxonomic assignment were combined with metadata using the phyloseq package (McMurdie and Holmes, 2013). ASVs classified as chloroplast, mitochondria, or eukaryota were removed from the dataset. Samples with 1,000 or less reads were discarded. After this filtering, 3,737 ASVs and 59 samples were used for statistical analysis using the microbiomeMarker and vegan packages (Cao et al., 2023; Oksanen et al., 2022). The percent relative abundance of each microbial phylum for each sample was calculated and ASVs with a total relative abundance <0.001% were removed. All additional analyses were conducted at the genus level. Plots were constructed using ggplot2 and tidyverse packages (Wickham et al., 2021, 2019). Alpha diversity was calculated using Shannon and Chao1 diversity indices. Beta diversity was estimated using the Aitchison distance and patterns in microbial community composition were visualized by Principal Component Analysis (PCA). Heatmaps were generated using the Ampvis2 (Andersen et al., 2018). To test for differences in beta-diversity between groups, an analysis of similarities (ANOSIM) within the vegan package was used using 999 permutations. Analysis of Compositions of Microbiomes with Bias Correction (ANCOMBC) were used to find significantly enriched and differentially abundant taxa between sites.

### 2.5 Temporal trends

#### 2.5.1 Field surveys

To study the effects of acidification on cover and necrosis of *P. ottomuelleriana,* transects of permanent photographic quadrats (25 x 25 cm) were performed at 2 m depth in both study sites over time. A total of 111 photographs (CO_2_ vent = 55; Ambient pH site = 56) were taken after summer from 2016 to 2020 (except spring of 2020 due to the COVID-19 lockdown). Cover (total area of living tissue) and tissue necrosis (dead tissue or denuded skeleton) was analyzed with the ImageJ image software (National Institutes of Health, https://imagej.net/ImageJ).

#### 2.5.2 Multilevel Bayesian models

To estimate the differences in the temporal trends of the total area and the necrosis area of *P. ottomuelleriana* between CO_2_ vent and Ambient pH site, we used multilevel generalized Bayesian models. We fitted two separate models, one with total area and another one with the necrosis as response variables. The fixed effects of both models were acidification (two levels, with and without acidification), mean temperature (°C), year and their interaction. We considered the ID of each quadrat as the random intercept. To account for the temporal non-independence of the total area and necrosis area data, we included an autoregressive-1 temporal process, which assumes neighbouring observations within the time series to be more similar than further ones. To see the prior information and the full model formula please refer to the SI Material and Methods.

Both models were fitted using the brms package v2.1.0 (Bürkner, 2018) in R v4.0.0 (R Core Team, 2020). Models were run for 10000 iterations, with a warmup of 1000 iterations.

#### 2.5.3 Multilevel Bayesian models diagnostics

To check the validity of our multilevel Bayesian models we ran a set of diagnostics. We inspected model convergence by visually examining trace plots and using Rhat values (the ratio of the effective sample size to the overall number of iterations, with values close to one indicating convergence). We also evaluated the model fit by exploring the distribution of the residuals, their variance, and the posterior predictive checks (Fig. S2-S4). All the diagnostics suggested a good model fit.

## 3. Results and Discussion

### 3.1 Differing compensatory responses to cope with low pH by adjusting skeletal properties

Our mineralogical and structural results reveal the compensatory responses to acidification may be more complex and distinct than initially expected. The encrusting skeletons of *P. ottomuelleriana* were bimineralic (calcite and aragonite) and high-Mg calcite (HMC, mean = 8.90 ± 0.71 mol% MgCO_3_ in calcite), being thus potentially soluble to acidification. At the CO_2_ vent, this species showed a decrease of Mg-calcite (8.33 ± 0.45 mol%; *t*-test vs 9.47 ± 0.32 mol%; p = 0.02), carbonate skeleton thickness and zooid size (Fig. 2; Table S3). *P. ottomuelleriana* thus appears to reduce the solubility of its skeleton, by incorporating lower Mg-calcite content, and reduce its investment in calcification and growth to counter acidification and/or lower Ωc and Ωa saturation states at the CO_2_ vent, consistent with previous results in other bryozoan species (Lombardi et al., 2011; Swezey et al., 2017). The production of weaker skeletons under low pH conditions may also contribute to increased damage susceptibility and thus compromise the survival of this species in the near-future.

**Fig. 2.**
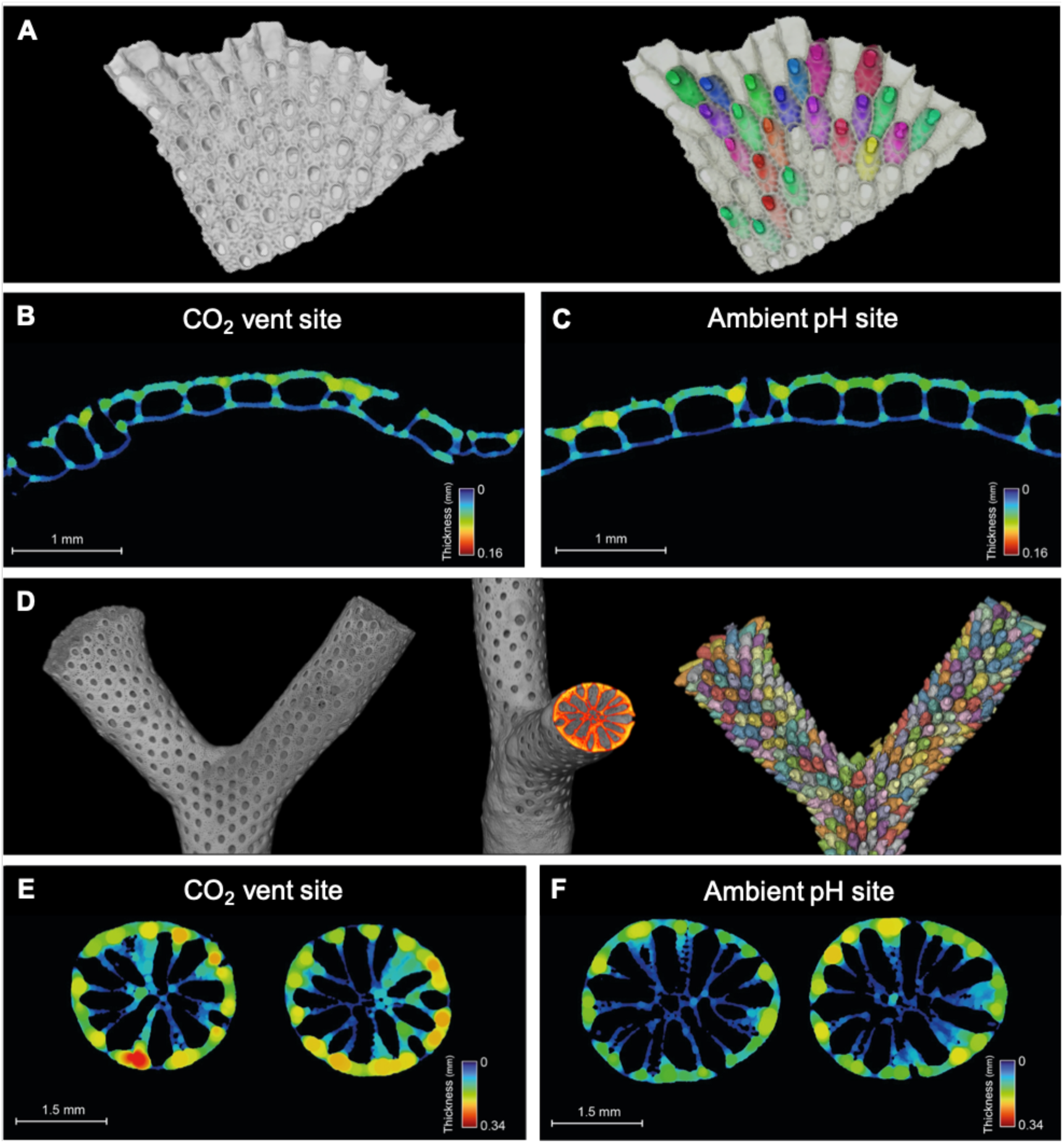
Micro-CT representative reconstruction of bryozoan species. A. Detail 3D image of carbonate skeleton of *Pentapora ottomuelleriana* (left) and zooid volumes reconstruction (right). B and C. 2D thickness map of *P. ottomuelleriana* from the CO_2_ vent and the Ambient pH site, respectively. D. Detail 3D images of carbonate skeleton of *Myriapora truncata* (left) with a distal 2D slice and zoid volumes reconstruction (right). E and F. 2D thickness map of *M. truncata* from the CO_2_ vent and the Ambient pH site.

In contrast, *M. truncata* had calcitic, IMC (5.03 ± 0.36 mol%) and thicker skeletons at the CO_2_ vent, likely due to the energy reallocation to calcification to overcome the decrease in pH (Fig. 2; Table S4). The higher investment of energy into calcification supports previous findings, where colonies of the coral species *Astroides calycularis*, that naturally co-occur at the studied sites, also experienced a shift to less porous and denser skeletons at the CO_2_ vent site (Teixidó et al., 2020). In addition, *Myriapora*’s skeleton is potentially less soluble (calcitic and IMC), than that of *P. ottomuelleriana*, which could confer to the erect bryozoan greater resistance to pH changes than the encrusting species and contributes to explaining why this species did not exhibit any significant mineralogical changes between sites. Organisms with slow calcification/growth rates, like *M. truncata*, may also have traits, such as lower metabolism, conferring them increased resilience to ocean acidification. For instance, this has already been observed in the scleractinian corals *Porites astreoides* and *Siderastrea siderea* (Martinez et al., 2019) and the red abalone *Haliotis rufescens* (Swezey et al., 2020). Supporting this explanation, *M. truncata* also exhibited a higher resistance to a wide range of thermal stress conditions than *P. fascialis*, which has faster growth rates (Pagès-Escolà et al., 2018).

### 3.2 Microbial communities seem to be stable and adapted to future pH conditions

This study is the first microbiome characterization of both bryozoan species and one of the first in this phylum (Boscaro et al., 2022; Bourne et al., 2013; Li et al., 2019; Patin et al., 2019). Therefore, we provide new insights on the identification of both bryozoan stable microbial associates and their potential functions within both holobionts. We also document novel patterns about microbiome shifts in response to future ocean acidification in bryozoans for the first time.

A total of 3,737 ASVs were retained in the dataset after quality filtering and discarding rare ASVs and 29 microbial phyla were identified. While significant differences in community alpha diversity were evident between the bryozoan microbiome and the surrounding seawater microbiome (p < 0.05), no significant differences were found between the bryozoan microbiome from both sites (two-sample Kolmogorov-Smirnov test; p > 0.05; Fig. S5). The effects of acidification thus seem not to affect the microbial alpha diversity in colonies of both bryozoan species. Other studies showed that acidification induced changes in microbial alpha diversity in coral species. These shifts in community composition are likely a result of reduced ability to regulate and/or exclude incoming microbial taxa from the surrounding environment (McDevitt-Irwin et al., 2017; Meron et al., 2011, p. 201; Morrow et al., 2015).

The beta-diversity of the bacterial community between species and sites was significantly different (ANOSIM; p < 0.05) but the PCA showed that there were more differences between the two species than between the two sites within the same species, thus highlighting the strong link between microbiome and holobiont (Fig. 3; Fig. S6). Microbiomes from both species also shifted in a similar way under future pH conditions (Fig. 3; Fig. S6). Our findings thus suggest that microbial communities of both bryozoan species are stable and adapted to future pH conditions. At Phylum level, the microbiome composition at both sites was mostly composed of Proteobacteria, Bacteroidota and Planctomycetota (Fig. S7 and S8). Previous studies also found Proteobacteria dominated microbiomes of the bryozoan species *Bugula neritina* (Li et al., 2019; Patin et al., 2019). Within the Proteobacteria, the microbiome from bryozoan colonies was dominated by the orders Cellvibrionales, Rhizobiales and Rhodobacterales, which are generally abundant in corals (McDevitt-Irwin et al., 2017). Abundant bacterial orders of the Bacteroidota included Flavobacteriales and Chitinophagales. Remarkably, Flavobacteriales are suggested to be opportunistic pathogens and important in coral diseases (Gignoux-Wolfsohn and Vollmer, 2015). Within the Planctomycetota, the microbiome was dominated by the order Pirellulales, which is composed by anaerobic ammonia-oxidizing bacteria and is also a dominant order in some coral species (Lawler et al., 2016). Members of this order also likely contribute to the removal of metabolic waste within the host (Mohamed et al., 2010).

**Fig. 3.**
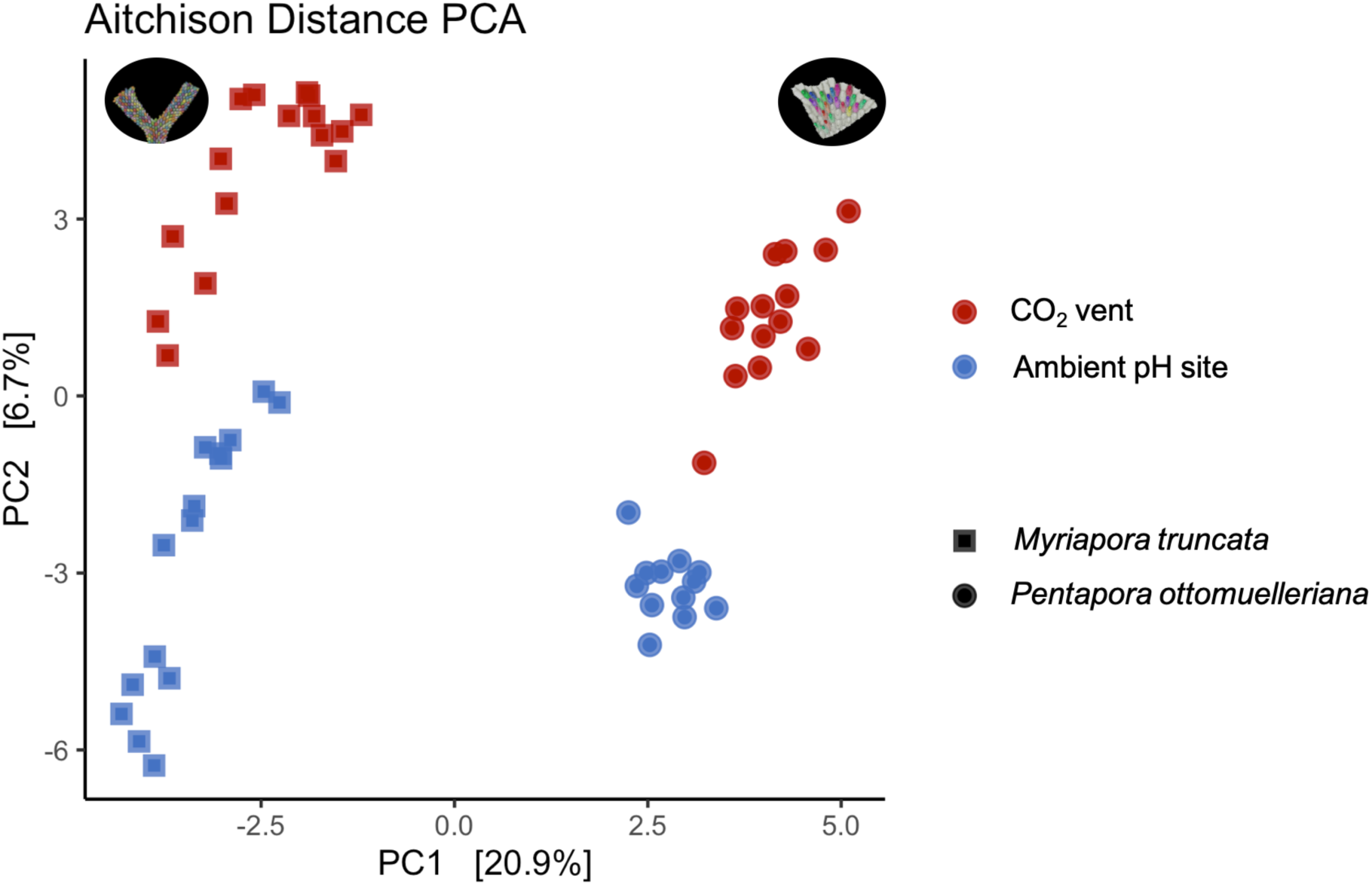
Principal component analysis (PCA) of the microbiome of both bryozoan species at the studied sites. Analysis of similarities (ANOSIM) was used to test for differences in beta-diversity between groups using Aitchison distance and 999 permutations. The beta-diversities of the bacterial communities between species and sites were significant (ANOSIM; P = 0.001).

Both bryozoan species shared nine abundant bacteria genera (BD1-7 clade, *Blastopirellula*, *Endozoicomonas*, *Arenicella*, *Aquimarina*, *Pseudahrensia*, *Croceitalea*, *Aquibacter* and *Subgroup 10*) at both sites, which indicates they are intimately related to the host and are probably part of their core microbiome (Fig. 4). Consistent with this, two of these dominant bacterial genera (BD1-7 clade and *Aquimarina*) and the genus *Mycoplasma*, which was only found in *M. truncata*, are suggested to be a key part of the microbiome of various temperate Gorgoniidae and are likely essential to the holobionts (van de Water et al., 2018; Van De Water et al., 2017). Interestingly, two dominant bacterial genera, “*Candidatus* Endobugula’’ and *Endozoicomonas*, of which the former was only observed in *P. ottomuelleriana*, have been previously retrieved from other bryozoan species (Li et al., 2019; Miller et al., 2016; Patin et al., 2019) (Fig. 4). Symbionts of “*Candidatus* Endobugula’’ have been isolated from several *Bugula* bryozoans and are known to be involved in the production of bryostatins, polyketides metabolites that provide chemical defense to the host (Davidson et al., 2001). In the case of *Endozoicomonas*, some members are commonly found in a range of marine organisms. These are suggested to have important roles in host health by providing nutrients through sulphur, nitrogen and/or methane cycling and synthesis of amino acids and by bacterial community regulation through the production of secondary metabolites (Morrow et al., 2015; Nishijima et al., 2013; Raina et al., 2009; van de Water et al., 2018).

**Fig. 4.**
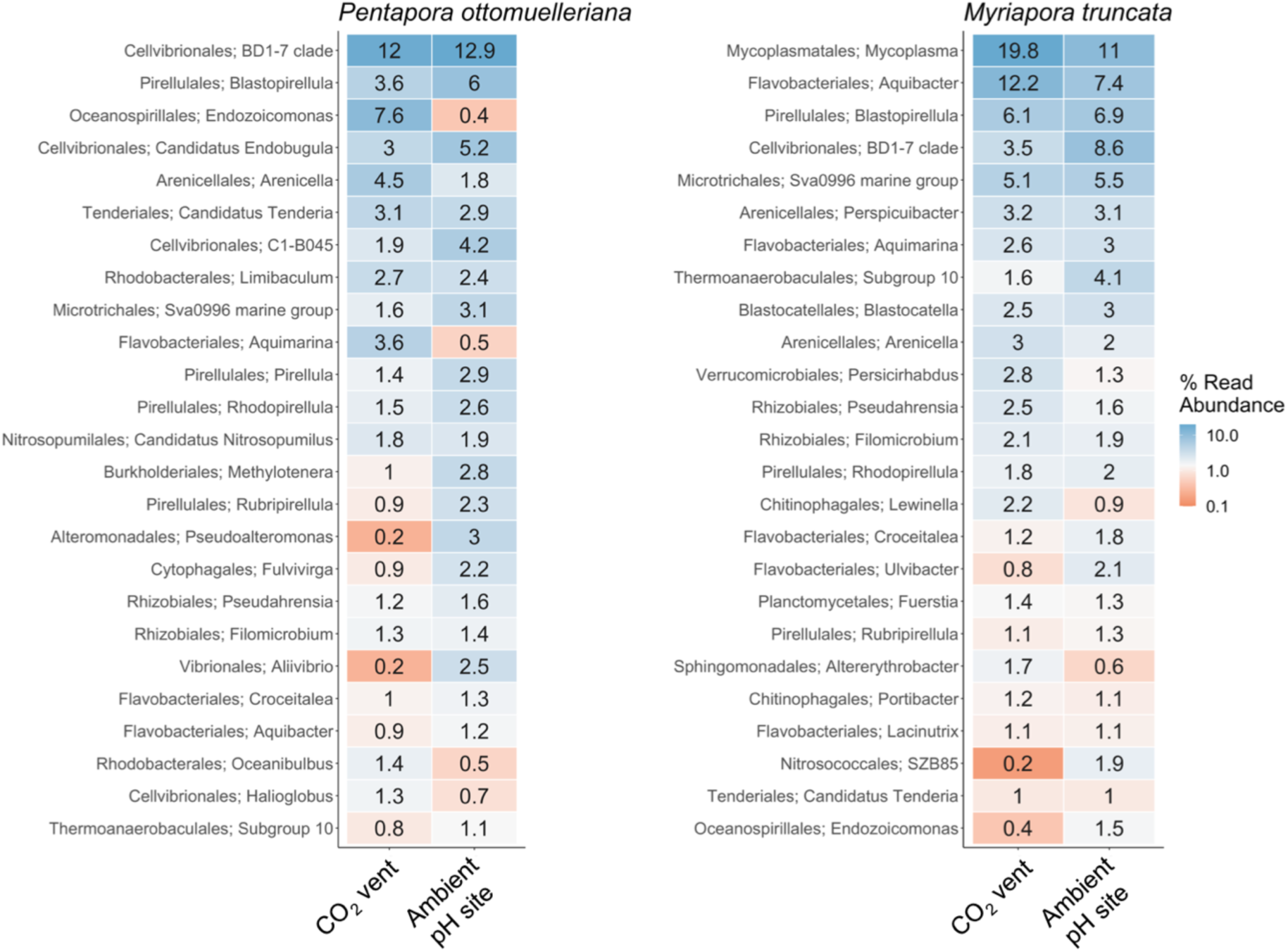
Relative abundance of bacterial genera in the microbiome of both bryozoan species at the studied sites. The color code indicates relative abundance, ranging from orange (low abundance) to dark blue (high abundance).

### 3.3 Microbial shifts under low pH conditions

While the bryozoan microbiomes were generally stable, a trend towards bacteria involved in organic matter degradation (*Lutibacter*, *Ulvibacter* and *Zobellia*; Flavobacteriales) were observed in both species under low pH conditions (Fig. 5). In particular, members of the genus *Zobellia* and *Ulvibacter* have specialized in the degradation of polysaccharides, a major component of marine organic matter (Nedashkovskaya et al., 2004; Salmeán et al., 2018), and *Lutibacter profundi* in degrading organic matter (Wissuwa et al., 2017). Consistent with these results, declines in pH increases the polysaccharide and other organic matter degradation (Piontek et al., 2010).

**Fig. 5.**
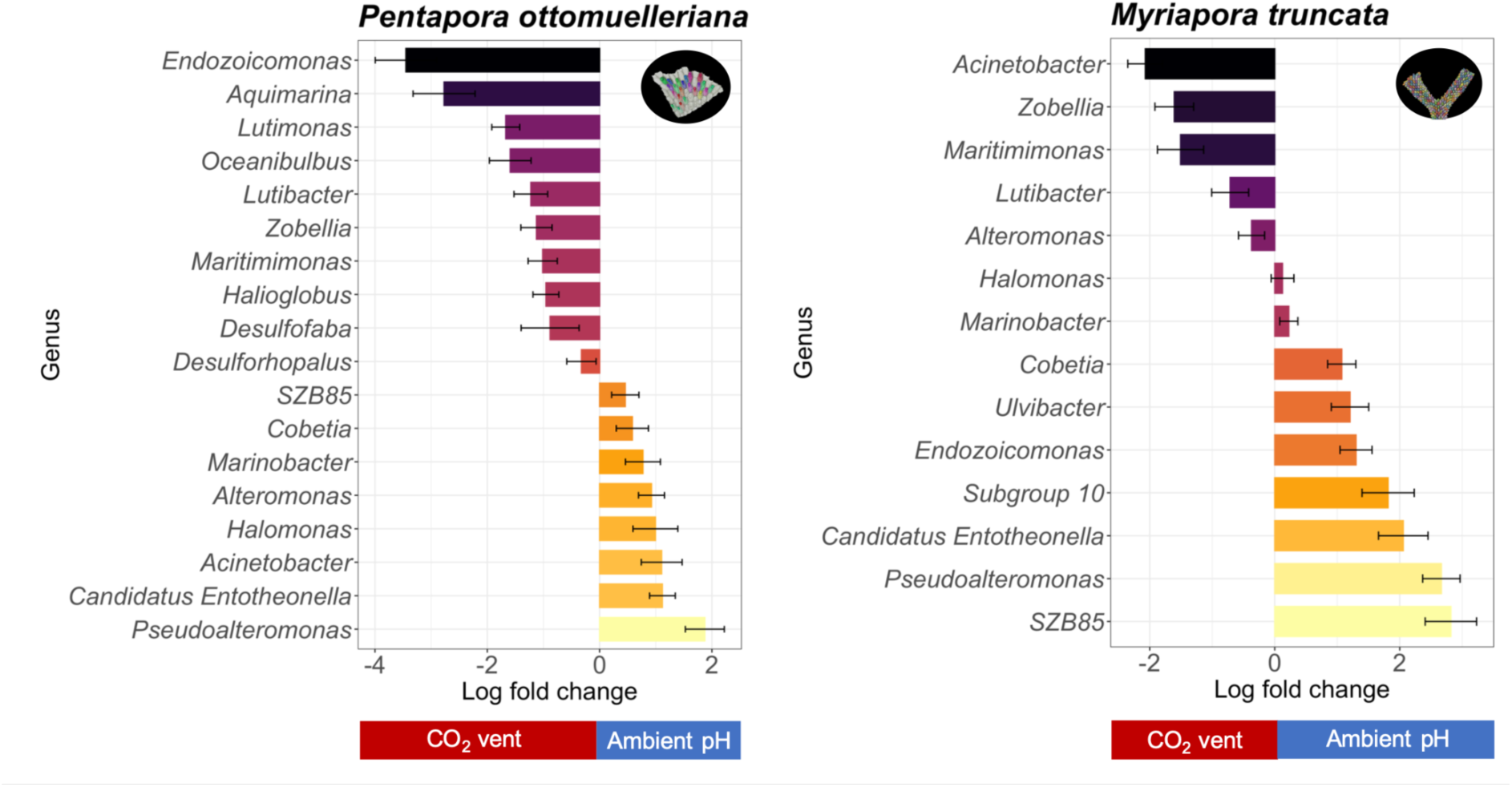
Abundance of bacterial genera in the microbiome of both bryozoan species that live in the CO_2_ vent site versus those that live in Ambient pH site according to ANCOMBC analysis. Natural log fold differences in abundance are shown with SD. A positive log fold change indicates a higher abundance in a bacterial genus in ambient pH conditions.

In *P. ottomuelleriana*, sulphate-reducing bacteria, strict anaerobes such as *Desulforhopalus* (Desulfobulbales) and *Desulfofaba* (Desulfobacterales) were enriched at the CO_2_ vent (Fig. 5). These bacteria have also been associated with higher rates of organic matter degradation, suggesting their role in carbon degradation (Tangherlini et al., 2021). *P. ottomuelleriana* encrusts the substrate forming an anaerobic environment which could thus favour the proliferation of anaerobic bacteria as suggested for corals (Yang et al., 2020). Our findings thus suggest that acidification modifies the microbiome environment of this bryozoan species, inducing a shift of the community towards anaerobes under these acidic conditions. This shift from aerobic to the anaerobic and facultative anaerobic bacteria under acidic conditions was previously found in other marine invertebrates such as the bivalve *Crassostrea gigas* (Zhang et al., 2022).

The increase of the relative abundance of sulphate-reducing bacteria may also be a mechanism to counteract the low pH as these bacteria may increase local alkalinity and induce CaCO_3_ precipitation (Cacchio et al., 2012; Gallagher et al., 2012). This could explain how *M. truncata* increases calcification under low pH conditions. In agreement, the extrapallial fluid of the eastern oyster *Crassostrea virginica*, which is responsible for shell formation, is enriched in sulphate-reducing and denitrifier bacteria (Sakowski et al., 2020).

Other genera involved in sulphur cycling showed opposite patterns such as *Alteromonas* (Alteromonadales), which was the dominant genus at the CO_2_ vent in *M. truncata* and was depleted in *P. ottomuelleriana,* or *Endozoicomonas* (Oceanospirillaes) which was enriched in *P. ottomuelleriana* and depleted in *M. truncata* at the CO_2_ vent. Similarly, species-specific changes in *Endozoicomonas* in response to low pH conditions were previously found in coral hosts, with a significant increase (Shore et al., 2021) or loss (Morrow et al., 2015; O’Brien et al., 2018; Shore et al., 2021; Webster et al., 2016). Shore et al. (2021) suggested the enrichment in *Endozoicomonas* may be beneficial to the host under ocean acidification conditions. By contrast, significant losses of *Endozoicomonas* may be lowering the host ability to control the growth of this bacteria (Shore et al., 2021).

### 3.4 Loss of bacteria with essential functions to the host and increase of bacteria previously associated with dysbiosis

We found that bacterial genera, which have essential functions to the host, were significantly depleted in both bryozoan species at the CO_2_ vent site, which suggest early warnings of potential deterioration of bryozoan health under near future ocean conditions. Some of the functions are biosynthesis of biologically active compounds with anti-bacterial, bacteriolytic, agarolytic and algicidal activities (*Pseudoalteromonas* and *Candidatus Entotheonella*), probiotic thermal protection (*Cobetia* and *Halomonas*) (Rosado et al., 2019), degradation of lipids and/or hydrocarbons (*Marinobacter*) (Mounier et al., 2014), and nitrification (*SZB85*), which might play key roles through facilitating the nitrogen cycle within cave ecosystems (Reboleira et al., 2022) (Fig. 5). These genera have been previously isolated from different coral, sponge and tunicate species, including bryozoans (Heindl et al., 2010; Pukall et al., 2001; Romanenko et al., 2005). The genus *subgroup 10* belonging to the phylum Acidobacteriota was also depleted in *M. truncata* at the CO_2_ vent and some members may be also beneficial to their host (e.g., production of diverse B-vitamins) as suggested in sponges (Engelberts et al., 2020; Webster et al., 2011).

An increase of bacteria previously associated with dysbiosis was also observed. In particular, *Acinetobacter*, which has commonly been isolated from marine invertebrates such as corals, was enriched in *M. truncata* and some members are pathogens (*Acinetobacter baumannii*) (Peleg et al., 2008) (Fig. 5). In *P. ottomuelleriana*, *Halioglobus* and *Oceanibulbus* also increased at the CO_2_ vent. *Halioglobus* has been found in bleached corals and may be associated to bacterial dysbiosis (Xu et al., 2023) and *Oceanibulbus* belongs to the disease coral-associated Rhodobacterales and some members of this order also increased with lower pH (Meron et al., 2011; Soffer et al., 2015; Sunagawa et al., 2009).

### 3.5 Ocean warming and acidification accelerated bryozoan mortality rates

We also observed potential health deterioration of bryozoan populations through time in the encrusting species. While our model suggests that the increasing seawater temperature, by itself, drove a decline in the total area in both studied sites, the combined effect of increasing seawater temperature and acidification accelerated this area loss in the CO_2_ vent (Fig. 6a,b; Table S1; Fig. S9). Similarly, our model suggests the necrosis increased at a faster pace in the CO_2_ vent site than in the Ambient pH site, likely due to the low pH (Fig. 6c,d; Table S2). However, its effects are uncertain, underscoring the need for caution when interpreting these data (Fig. 6c,d).

**Fig. 6.**
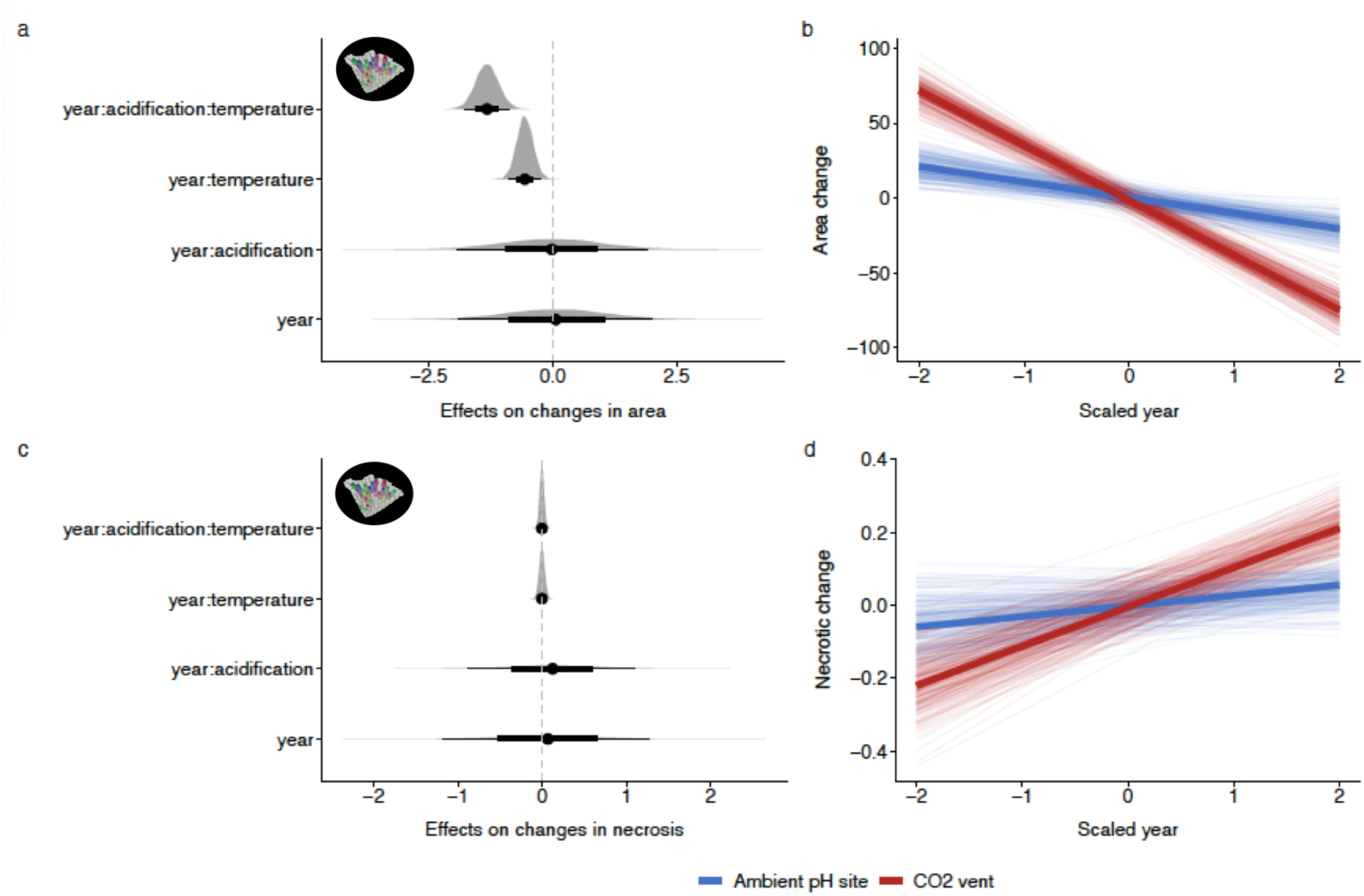
Effects of acidification and mean temperature on the total area and necrosis of *Pentapora ottomuelleriana*. (a,c) Effect of year, mean temperature, acidification and their interaction to the (a) scaled value of total area (cm^2^) and (c) necrosis. The reported values are the highest posterior density median values (circles), with 50% (thickest bars), 80% and 95% (thinnest bars) uncertainty intervals. (b,d) Represent the temporal trend of the (b) scaled area and (d) necrosis in the CO**_2_** vent and Ambient pH site. The lines represent the predictions from the multilevel Bayesian models, where thin lines correspond to the predictions drawn from each of the 250 posterior samples of the model. Thick lines represent the mean outcome of the model.

Consistent with our results where we found that the increasing seawater temperature, alone, was already a driver of loss of bryozoan cover, the 2015–2019 period was the warmest on record in the Mediterranean Sea (Garrabou et al., 2022). During this period, several mass mortality events of a range of marine taxa were reported, being bryozoans the second most affected taxonomic group (Garrabou et al., 2022, 2019). Specifically, a congeneric species of *P. ottomuelleriana* (*P. fascialis*) and other encrusting bryozoan species (*e.g., Schizobrachiela sanguinea*) were affected (Garrabou et al., 2022, 2019; Gómez-Gras et al., 2021; Pagès-Escolà et al., 2018). Our findings thus suggest that while ocean warming is already posing a threat for this bryozoan species, ocean warming effects will likely be enhanced by the projected pH decline by the end of the century. These results emphasize that responses cannot be predicted from the analysis of individual-stressor impacts alone and highlight the importance of considering these two major stressors in assessing future potential responses.

## 4. Conclusion

Our results highlight novel, differing and unpredictable responses on how bryozoans can cope with low pH. While both bryozoan species show indicators of acclimatization by having compensatory mechanisms (adjusting skeletal properties) and stable microbial communities under acidification conditions, the bryozoan status is still dependent on the effects of warming and other local environmental impacts. Consistent with this, we found the most adverse impacts arises from the combined effects of warming and acidification in the encrusting species *P. ottomuelleriana*. These findings thus suggest that *P. ottomuelleriana* may be particularly at risk to future impacts of warming and acidification in the Mediterranean Sea. The slow-growing *M. truncata* species is apparently more resilient to ocean acidification although the observed microbiome shifts may indicate a decrease of its resilience to near future ocean conditions. Both bryozoan species could thus potentially act as sentinels providing early warnings of health deterioration in bryozoan populations. Our results thus emphasize that more multidisciplinary studies on long-term impacts using both stressors, multi-species, and their microbiome are needed to better predict how global environmental change will impact bryozoans.

## Acknowledgments

We thank C. Paradela for her support in the acquisition of the microCT scans, S. Álvarez and J. Ivañez for their support with the XRD, CORELAB laboratory (Universitat de Barcelona) for his support to analyze micro-CT data and A.M. Bonacolta for his support with the microbiome analyses.

