## Supplementary Material for "Long-term effects of ocean acidification and its interaction with warming on calcifying organisms and their associated microbiome: bryozoans as emerging sentinels of global change?"

**This file contains:**

SI Materials and Methods

Figures S1-S9

**SI Material and Methods**

**X-ray computed tomography**

The initial step involved applying a non-local means algorithm to denoise the scalar volume data, followed by implementing a delineate filter to enhance the edges of objects, thereby improving object differentiation. The second step was segmented the image data to divide the bryozoan images into various subregions (i.e. carbonate skeleton and zooids volumes) which was obtained through the application of multiple morphological transformations. Finally, the subregion analysis allowed the extraction of various measurements from the images, such as volume, length, and surface area. Additionally, a thickness of carbonate skeleton analysis was measured using the average object thickness module of Avizo®, which is given by a parallel plate model:

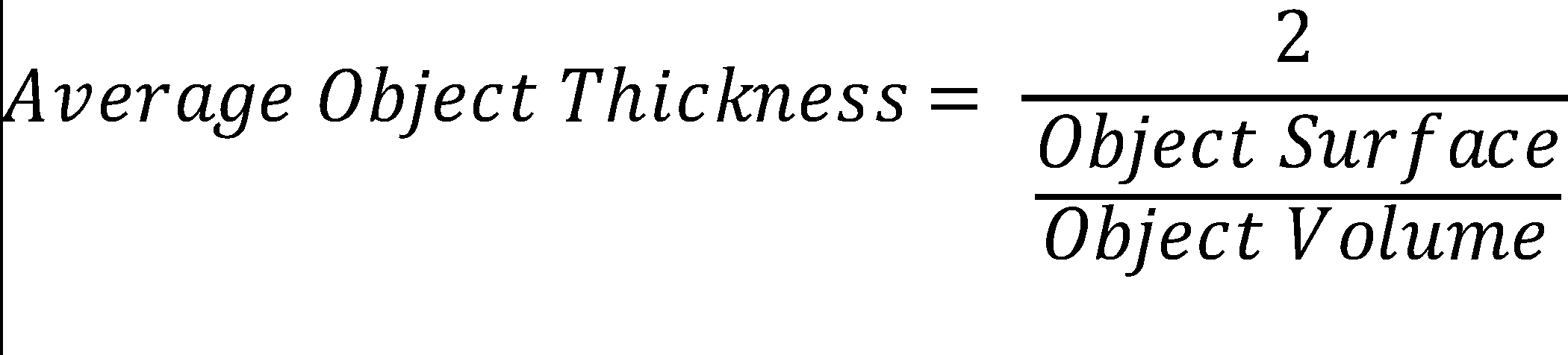

**Multilevel Bayesian models**

The structure of the model was:

${Y_{i}\sim Normal(\mu_{i}, \sigma)}$, eq1

$\mu_{i}=\alpha+\alpha_{j[i]}+\beta_{1}A_{[i]}+\beta_{2}{MT}_{[i]}+\beta_{3} {Year}_{[i]}+\beta_{123} A,MT,{Year}_{[i]}+AR(1)+\epsilon_{ij}$, eq2

where $Y_{i}$represents the total area or necrotic tissue of *P.* *ottomuelleriana* of the *j*th quadrat ID and is given by a Normal distribution with mean *µ* and a variance *σ*. $\alpha$represents the intercept and $\alpha_{j}$is the independent random intercept for quadrat ID (*j*). $\beta_{1}A$ is the effect of the discrete factor *acidification,* $\beta_{2}MT$ is the effect of the continuous factor *mean temperature,* $\beta_{3}Year$ is the effect of the continues factor *year*, and $\beta_{123}A,MT,Year$ is the interaction among these factors. *AR(1)* represents the autoregressive process of order 1.

We used weakly informed priors:

${\alpha\sim Normal(0,10)}$, eq3

${\alpha_{j}\sim Normal(\underline{\alpha}, \sigma_{\alpha})}$, eq4

$\underline{\alpha}\sim Normal(0,10)$, eq5

$\beta_{1}\sim Normal(0,10)$, eq6

$\beta_{2}\sim Normal(0,10)$, eq7

$\beta_{3}\sim Normal(0,10)$, eq8

$\beta_{123}\sim Normal(0,10)$, eq9

$AR(1)\sim Beta(2,2)$, eq10

$\sigma\sim Exponential(1)$, eq11

$\sigma_{\alpha}\sim Exponential(1)$, eq12

**Fig. S1.** Temperature time series data in the CO_2_ Vent (Grotta del Mago; A) and the ambient pH site (Punta Vico, B). Daily mean temperature calculated from hourly measurements taken by in situ hobo sensors from 2016 to 2021 at 2 m depth. Temperature followed ambient seasonal fluctuations at both studied sites.

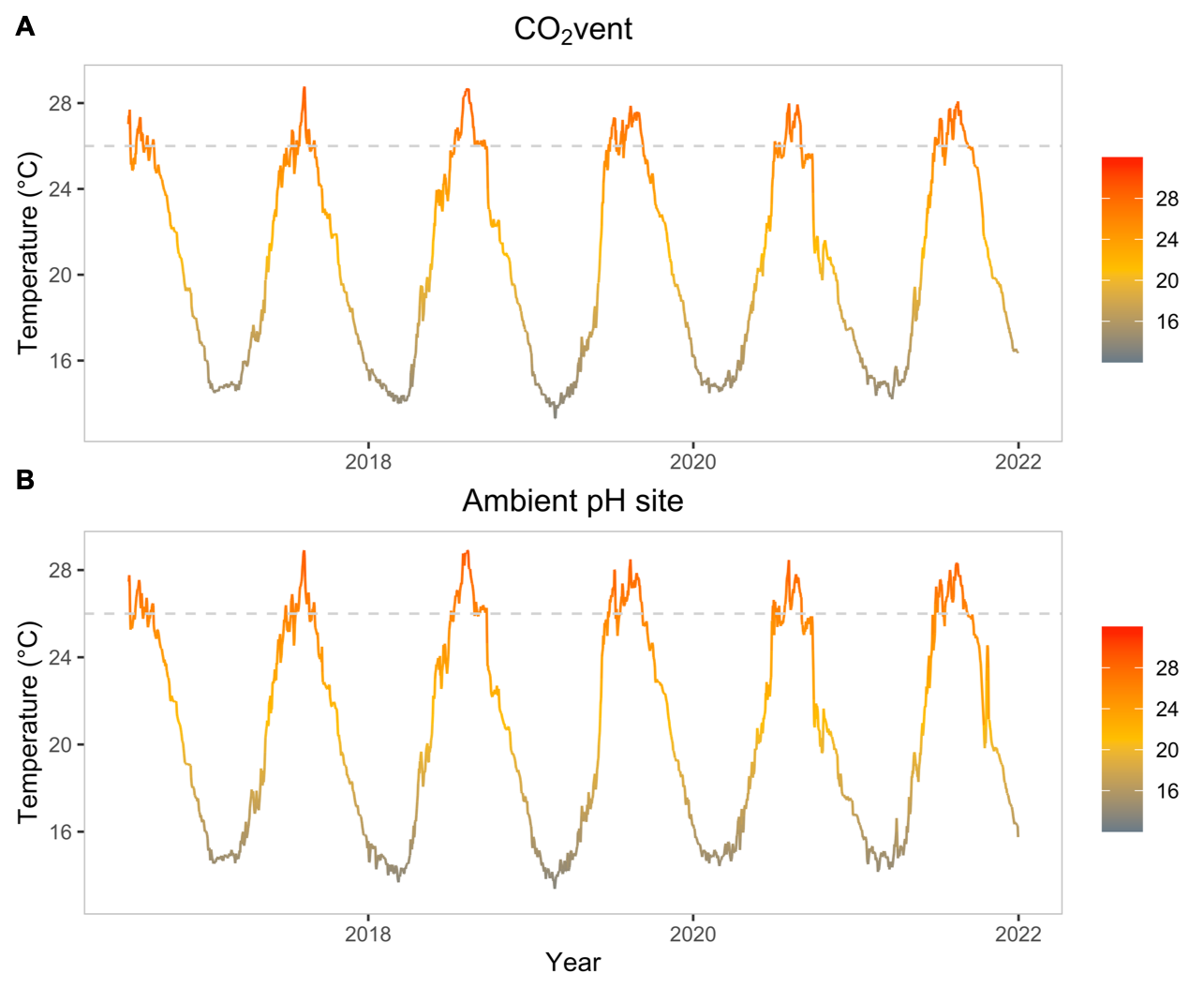

**Fig. S2. Distribution of the standardised residuals for the multilevel Bayesian models.** Residuals were examined for the (a) total area and (b) necrotic models.

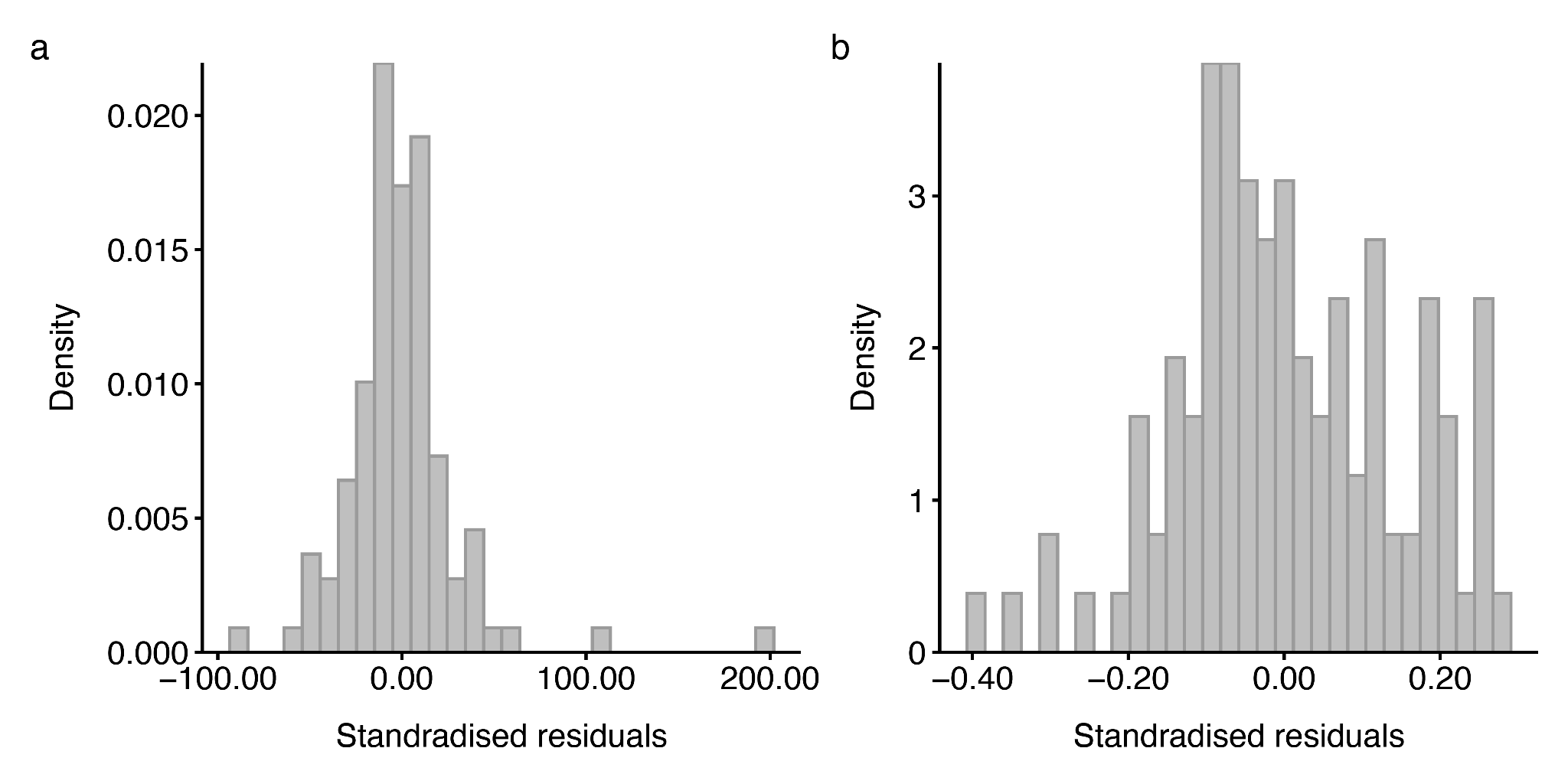

**Fig. S3. The standardised residuals vs the predicted values for the multilevel Bayesian models.** The standardised residuals vs the predicted values have similar variances for all models, suggesting that the equal variance assumption is met. Residuals were examined for the (a) total area and (b) necrotic models.

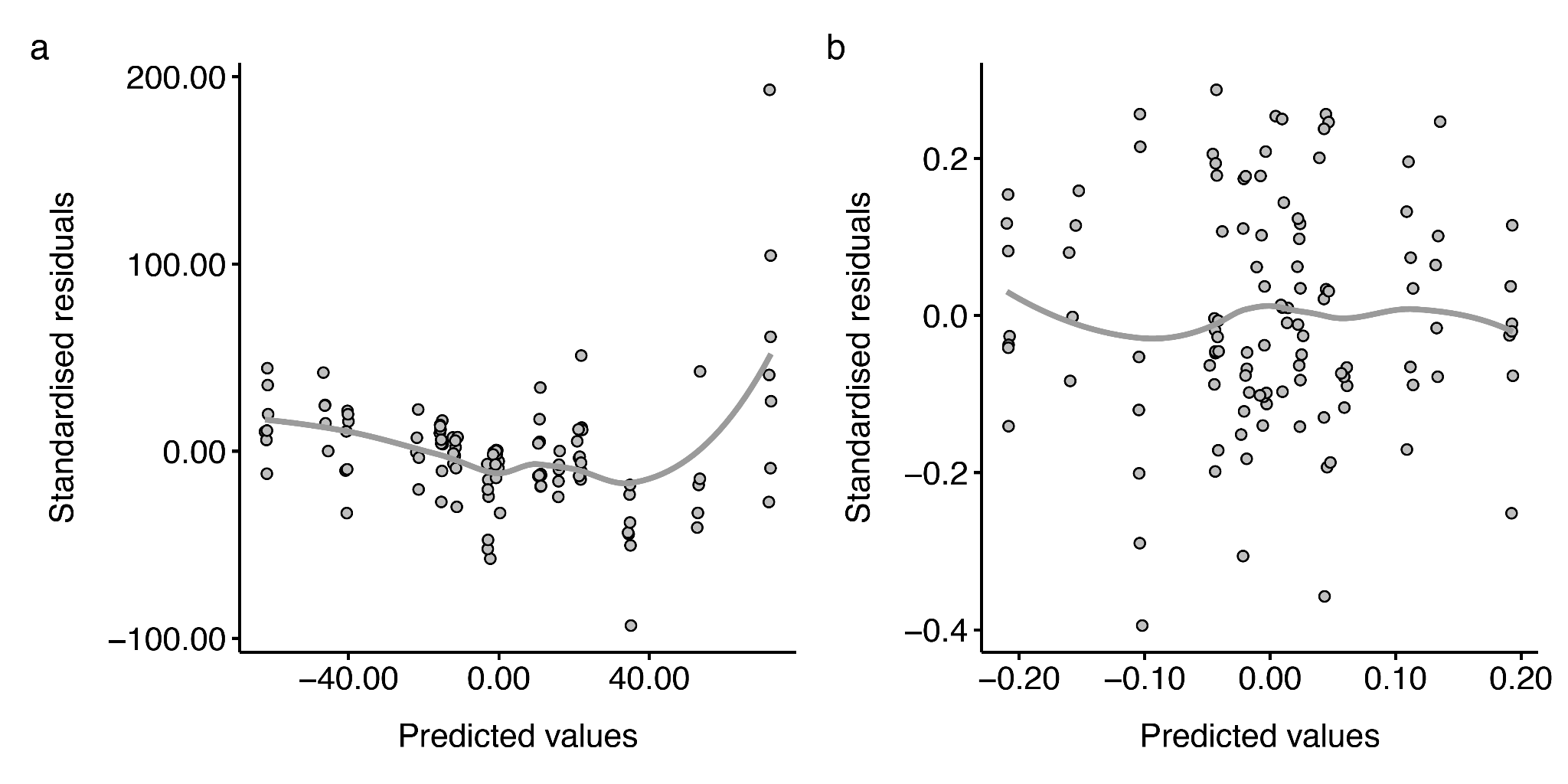

**Fig. S4.** **The posterior predictive checks for the multilevel Bayesian models.** The posterior predictive checks do not show strong discrepancies between our data (dark lines, y) from the predictions from the model (light grey lines, yrep) for any of the models. However, the model shows a slight underestimation of the true zero values. The posterior predictive checks were examined for the (a) total area and (b) necrotic models.

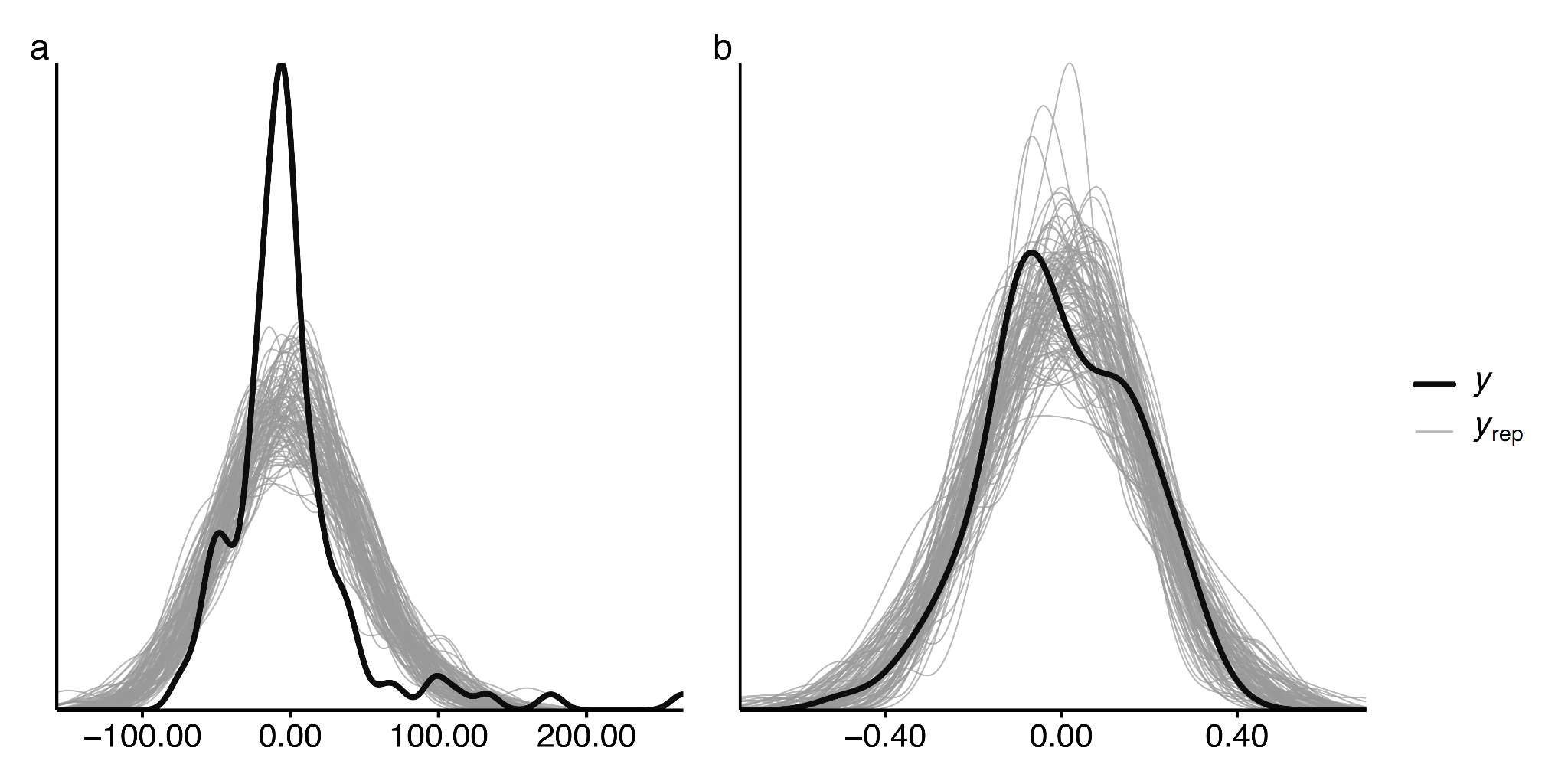

**Fig. S5. Alpha-diversity of the microbiome of both bryozoan species at the studied sites.** The alpha-diversities of the bacterial communities between both sites were not significant (two-sample Kolmogorov-Smirnov test; p > 0.05).

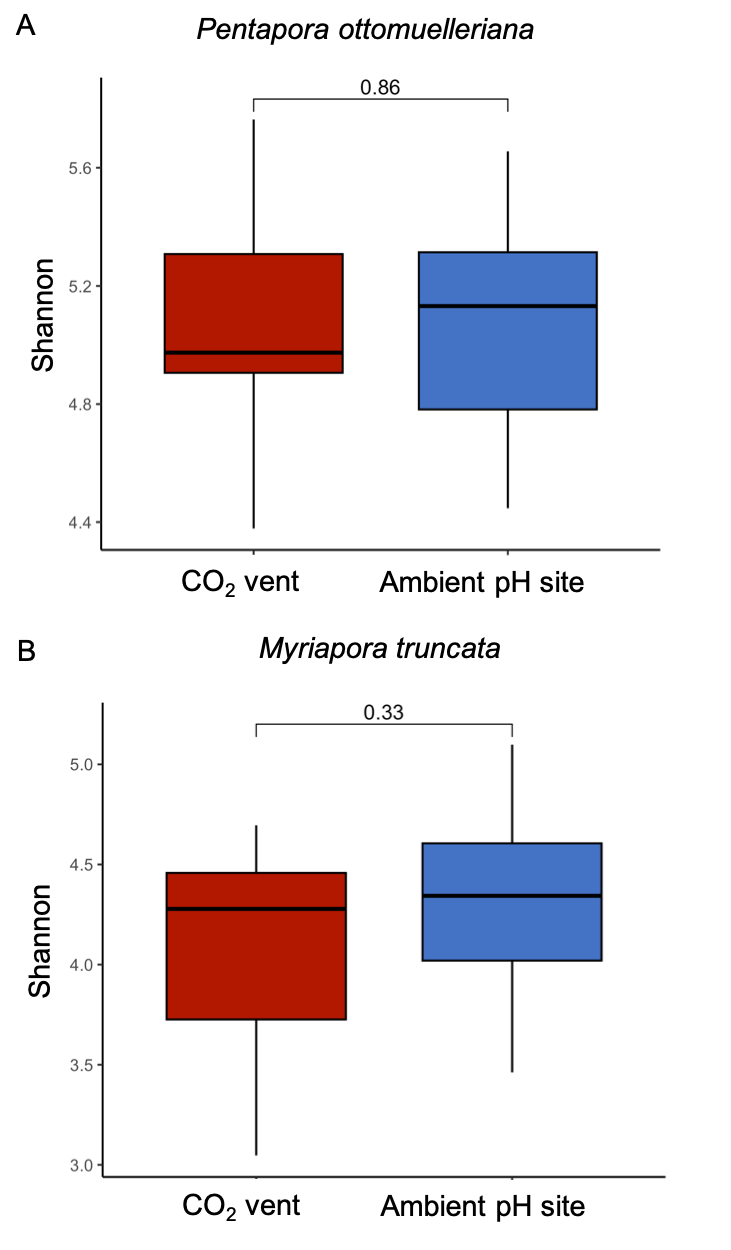

**Fig. S6. Beta-diversity of the microbiome of both bryozoan species and seawater at the studied sites.** Analysis of similarities (ANOSIM) was used to test for differences in beta-diversity between groups using Aitchison distance and 999 permutations. The beta-diversities of the bacterial communities between species and sites and seawater were significant (ANOSIM; P = 0.001).

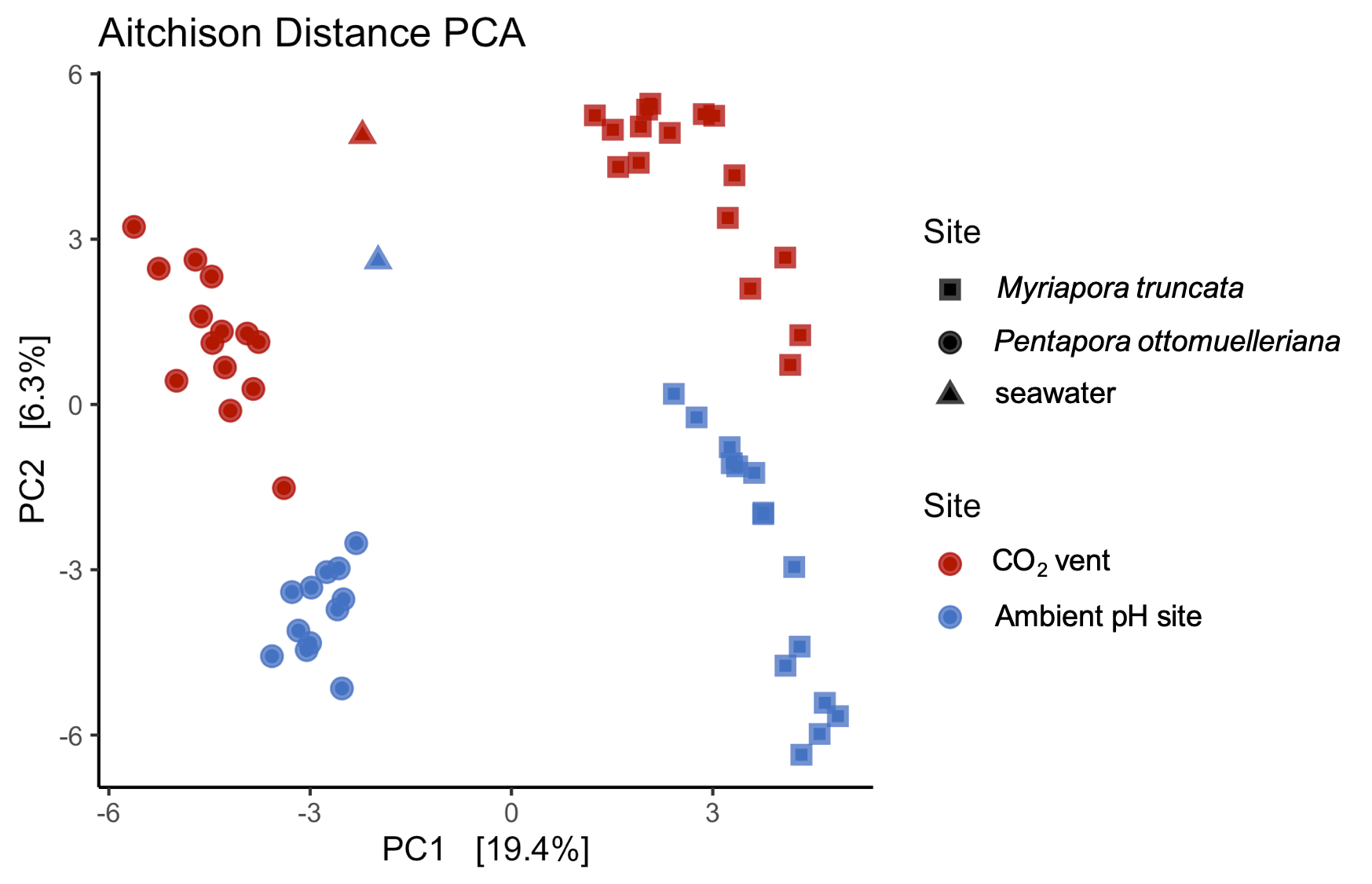

**Fig. S7. Bubble plot showing the relative abundance of the most abundant orders (%) associated with *Pentapora ottomuelleriana* at each site.** Circle sizes are proportional to the relative abundance and colored according to their phyla and are shown with SD. Only relative abundances greater than 1% are shown. “Other” represents all that could not be classified below Kingdom level.

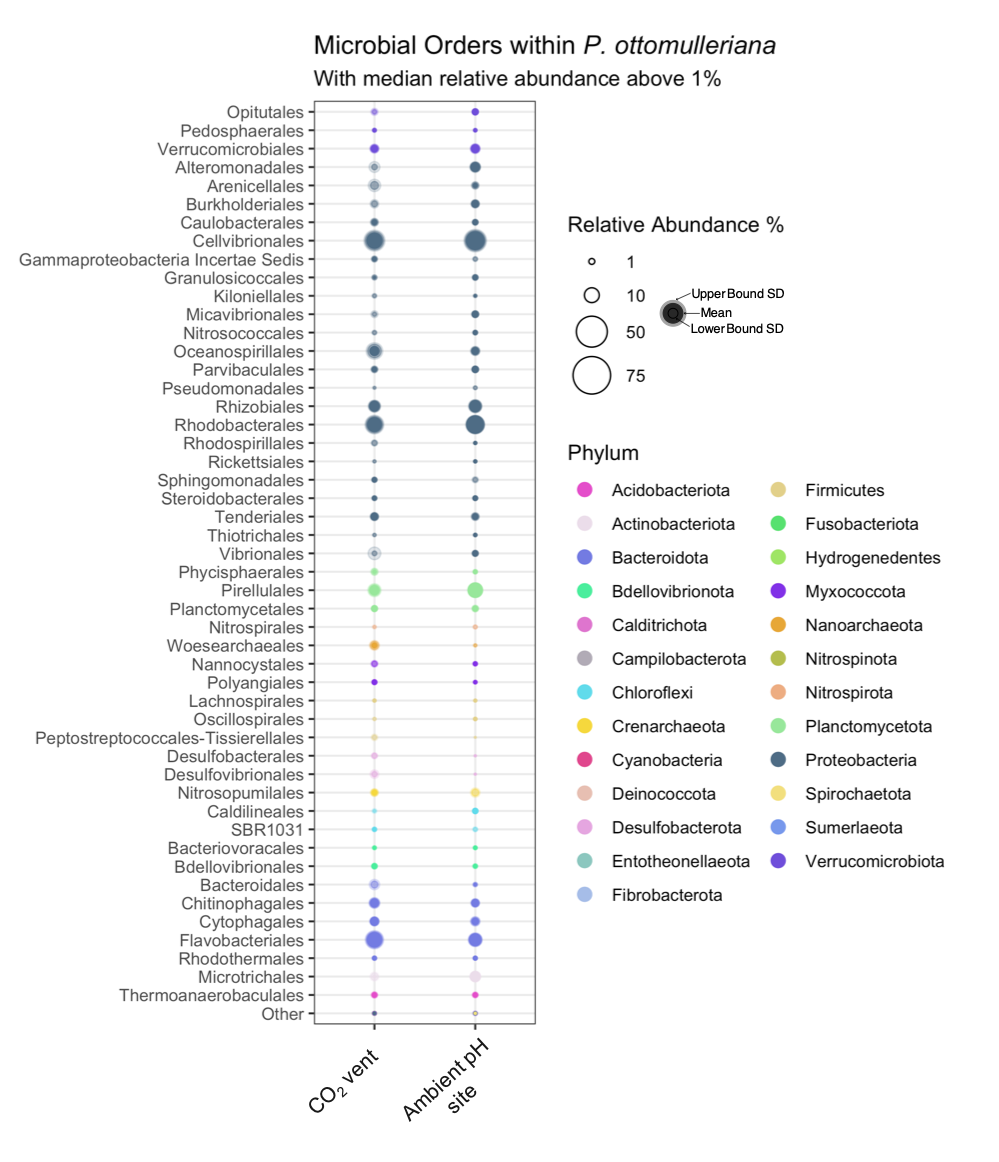

**Fig. S8. Bubble plot showing the relative abundance of the most abundant orders (%) associated with *Myriapora truncata* at each site.** Circle sizes are proportional to the relative abundance and colored according to their phyla and are shown with SD. Only relative abundances greater than 1% are shown. “Other” represents all that could not be classified below Kingdom level.

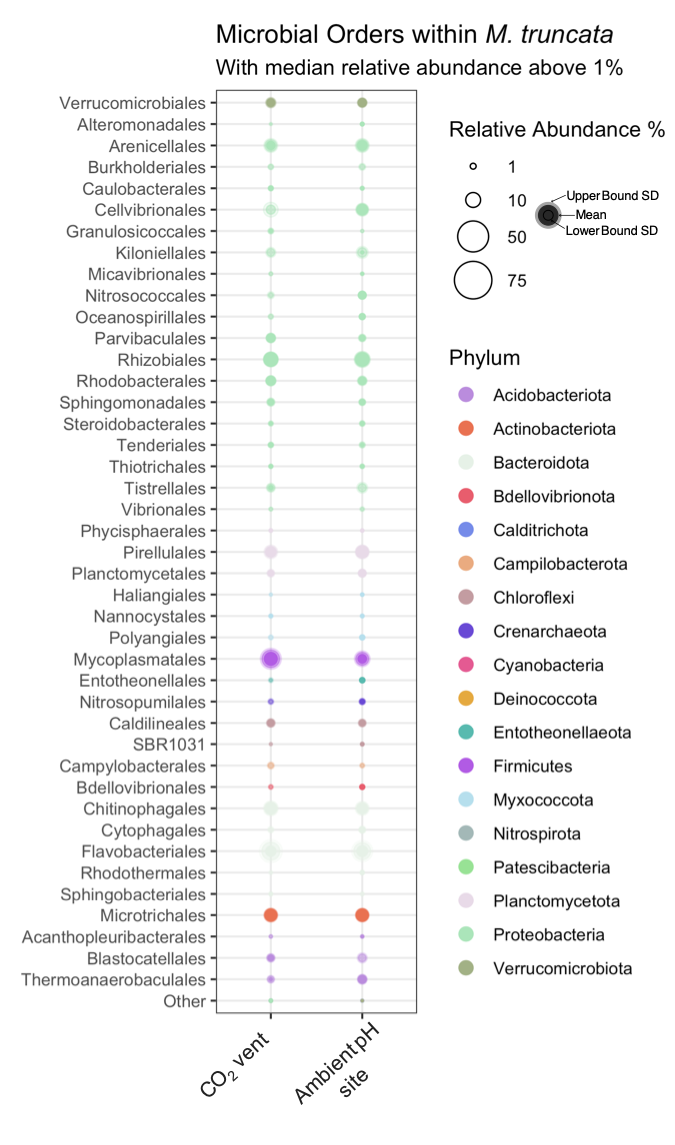

**Fig. S9. Images scaled with a 25 × 25 cm quadrat.** A decline and loss of the encrusting bryozoan *Pentapora* *ottomulleriana* (orange encrusting patches) is clearly observed from 2016 to 2019.

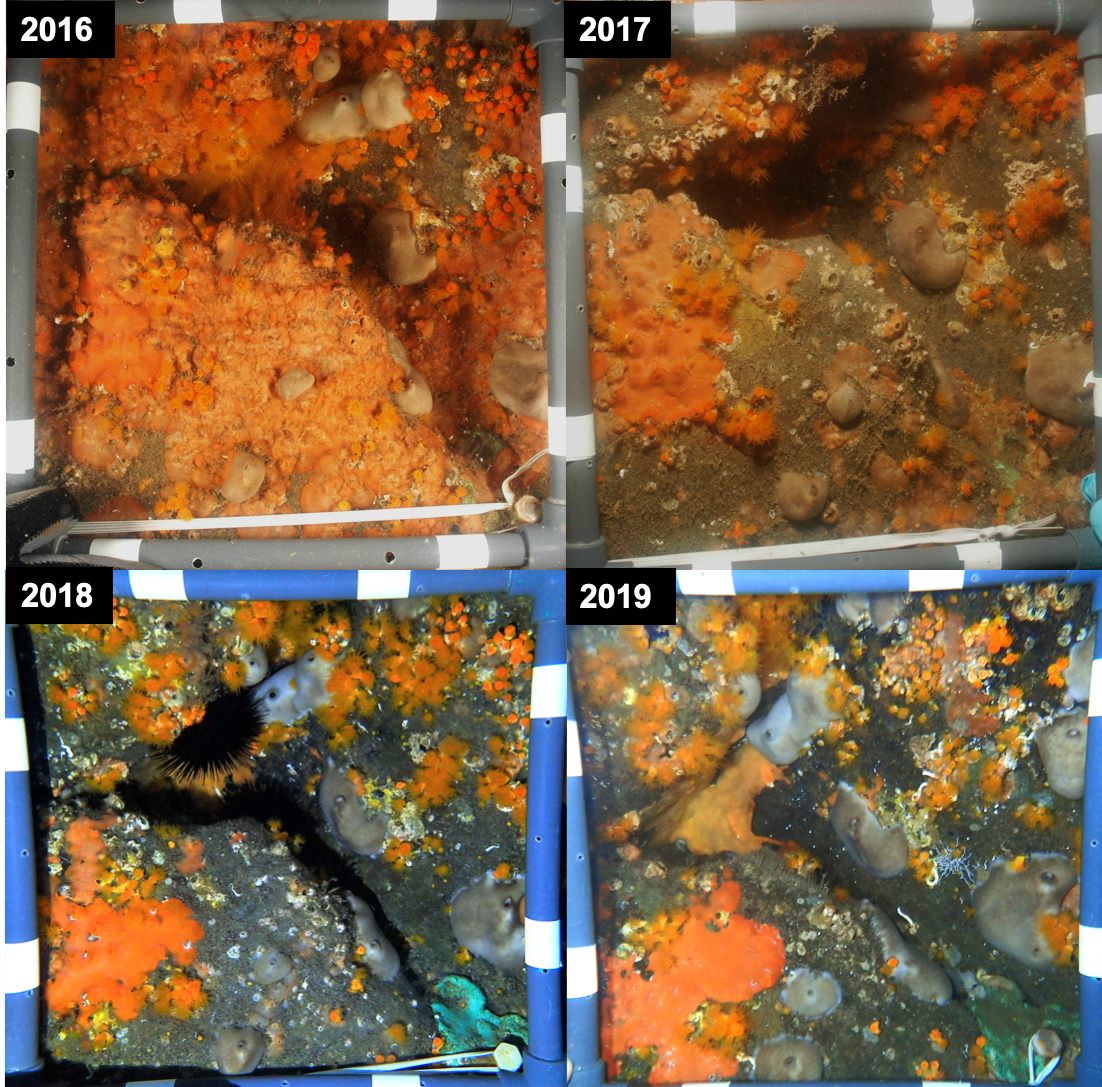
